# Structural analysis of the active site and DNA binding of human cytidine deaminase APOBEC3B

**DOI:** 10.1101/304170

**Authors:** Shurong Hou, Tania V. Slivas, Florian Leidner, Ellen A. Nalivaika, Hiroshi Matsuo, Nese Kurt Yilmaz, Celia A. Schiffer

## Abstract

APOBEC3s proteins (A3s), a family of human cytidine deaminases, protect the host cell from endogenous retro-elements and exogenous viral infections by introducing hypermutations. However, the ability to mutate genomic DNA makes A3s a potential cancer source. Of the 7 human A3s, A3B has been implicated as an endogenous cause for multiple cancers. Despite overall similarity, A3s have distinct deamination activity with A3B among the least catalytically active. Over the past few years, several structures of apo as well as DNA-bound A3 proteins have been determined. These structures revealed the molecular determinants of nucleotide specificity and the importance of the loops around the active site in DNA binding. However, for A3B, the structural basis for regulation of deamination activity and the role of active site loops in coordinating DNA had remained unknown. In this study, using a combination of advanced molecular modelling followed by experimental mutational analysis and dynamics simulations, we investigated molecular mechanism of A3B regulating activity and DNA binding. We identified a unique auto-inhibited conformation of A3B that restricts access and binding of DNA to the active site, mainly due to the extra PLV residues in loop 1. We modelled DNA binding to fully native A3B and found that Arg211 in the arginine patch of loop1 is the gatekeeper while Arg212 stabilizes the bound DNA. This model also identified the critical residues for substrate specificity, especially at the -1 position. Our results reveal the structural basis for relatively lower catalytic activity of A3B and provide opportunities for rational design of inhibitors that specifically target A3B to benefit cancer therapeutics.

## Introduction

APOBEC3s (A3s) are a family of cytidine deaminases that catalyse a zinc-dependent cytidine to uridine reaction on single strand DNA (ssDNA) or single strand RNA (ssRNA) (1–4). The family comprises of seven members that have either one (A3A, A3C, A3H) or two (A3B, A3D, A3F, A3G) zinc-binding domain (5). The two-domain A3s have a pseudo-catalytic N-terminal domain (NTD) and a catalytically active C-terminal domain (CTD). A3s play a key role in innate immunity by protecting the genome from exogenous and endogenous retro-elements through introducing G to A hypermutations (6–11). However, when overexpressed, their mutagenic activity can also cause modification of genomic DNA and thus promote tumorigenesis (12–14).

A3B has been identified as a significant enzymatic source of mutagenesis in a variety of cancers (15). Endogenous A3B is involved in the restriction of retro-element LINE-1 (16) and HBV (17,18). However when overexpressed, A3B can mutate the host genome to trigger cancer phenotypes (12). The up-regulation of A3B in tumours is correlated with both dispersed and clustered high occurrence of cytidine mutations, TP53 (tumour protein 53) inactivation, and poor patient outcome in cancer treatment (12,19–22). In addition, the genomic mutations preferentially occur at 5-TCA, 5-TCG, and 5-TCT trinucleotide motifs, which resemble the substrate preference of A3B in biochemical assays (19,21). Unlike other cancer sources, A3B executes an active mutational process, which means that a growing number of DNA mutations will be created. This will further benefit cancer evolution, for example, to help escape immune monitoring, outgrow, metastasize, and potentially acquire resistance to therapeutic treatments (23). Hence, in addition to its non-essential nature (24), A3B represents a promising target for novel anticancer drug development.

Over the past several years, crystal and NMR structures of human A3s (A3A, A3C; CTDs of A3B, A3F, A3G) in the apo state have been determined by our group (25–31) and others (32–45). In general, A3 proteins are structurally highly similar despite their distinct deamination activities. Among all human A3s, A3B-CTD and A3A share the highest sequence identity (Figure 1 and **Supplementary Figure S1A**); however, A3A is about 15-fold more active compared to A3B-CTD (44). The overall A3 structure consists of six alpha-helices and five beta-strands with the zinc-binding domain in the middle. Recently, our laboratory (46), along with two other groups, has determined the crystal structures of three A3-DNA complexes (A3A-DNA, chimeric A3B-DNA and rA3G-DNA)(45,47). When A3A binds to DNA, two major changes occur at the active site involving the side chain of Tyr132, which stacks against the DNA, and the gatekeeper His29, which locks the DNA in the active site. To facilitate crystallization of DNA-bound A3B, loop 1 of A3A was swapped into A3B to determine the structure. However, differences in loop 1 between the two A3s are mainly responsible for the difference in catalytic activity, as swapping loop 1 of A3B by that of A3A increases A3B activity by 10-fold (44). Loop 1 also exhibits the largest amino acid sequence difference between A3A and A3B (Figure 1). Loop 1 is longer in A3B with three more residues _206_PLV_208_. In addition, the DNA gatekeeper residue His29 in A3A (46), is missing in A3B and is likely replaced by one of the arginines within the unique triple arginine patch _210_RRR_212_. However, the role of loop 1 in regulating A3B’s catalytic activity and in coordinating DNA could not be determined by crystal structures.

**Figure 1.**
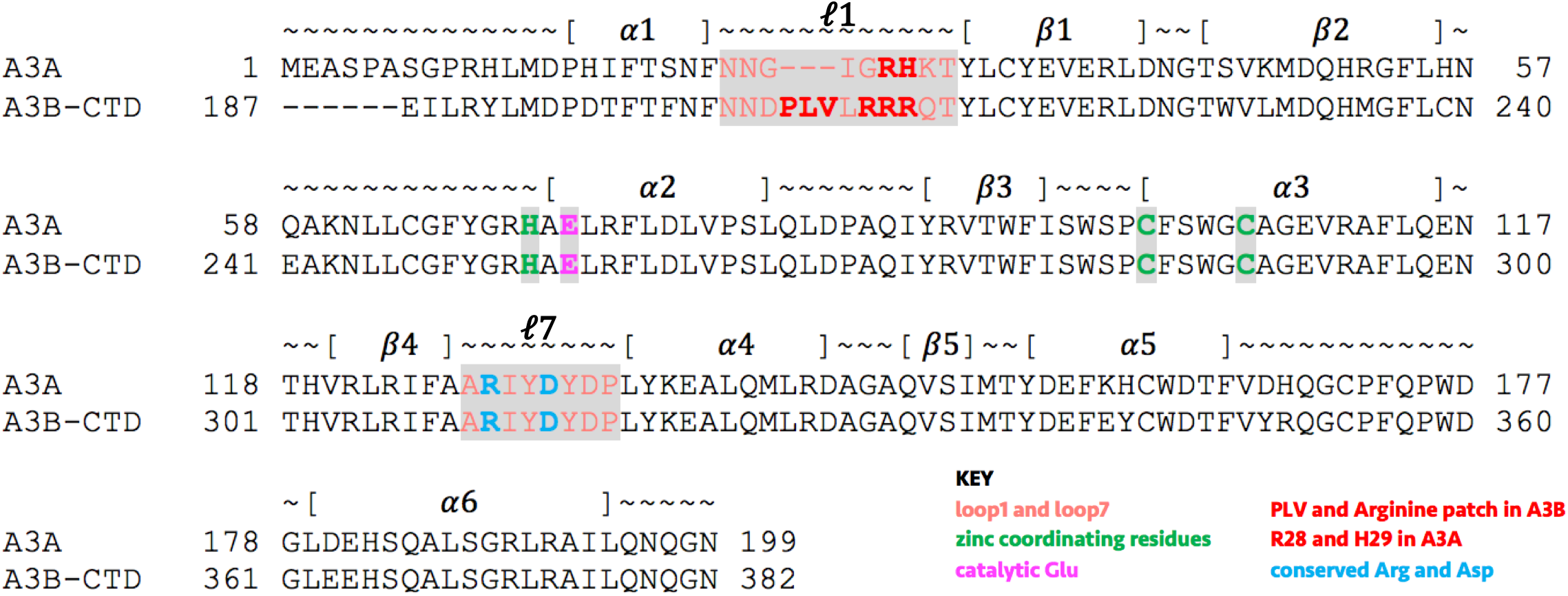
Amino acid sequence alignment of A3A and the catalytic domain of A3B.

To elucidate the molecular mechanism of how A3B structurally regulates its catalytic activity and DNA binding, we used a combination of molecular modelling with molecular dynamics simulations and experimental mutational analysis followed by fluorescence-anisotropy based DNA binding assays. We identified the key role of loop 1 in down-regulating A3B activity, and in binding DNA. A3B has an auto-inhibited conformation that is unique among human A3s, resulting from differences in loop 1 length and sequence, which explains the relatively low catalytic activity of A3B. We also present a structural model of the A3B-DNA complex that elucidates the molecular mechanism and determinants of DNA binding to A3B. The model and mutational verification identified Arg211 as the gatekeeper for locking DNA into the active site, which is further stabilized by Arg212. Overall, our results shed light into the structural regulation of A3 activity and differences in loop 1 coordination around the bound DNA, which may potentially lead to discovering anti-cancer drugs to benefit cancer therapeutics.

## Material and Methods

### Molecular modelling

A3B-CTD wild type apo structure was modelled based on human A3B-CTD isoform A sequence and A3B-CTD crystal structure (PDB: 5CQH) through program Modeller 9.15 using basic modelling. All DNA-bound structures were modelled based on both the apo crystal structure (PDB: 5CQH for A3B; PDB: 4XXO for A3A) and A3A-DNA co-crystal structure (PDB: 5KEG) through program Modeller 9.15 using advanced modelling. The DNA molecule of the bound models and A3A-DNA co-crystal structures were modified through program Coot to the oligo sequence (AATCGAA) that was used in the fluorescence anisotropy-based binding assay. The phosphate groups of 5’ A base were removed to prevent strong electronegative environment. AACCGAA was modelled similarly to test the molecular mechanism of substrate preference at -1 position. All DNA-bound structure models were then energy minimized through Protein Preparation Wizard from Schrodinger using default settings.

### Molecular dynamics simulations

All molecular dynamics simulations were performed using Desmond (48) from Schrodinger. The models were first optimized using Protein Preparation Wizard. The simulation systems were then built through Desmond System Setup using OPLS3 force field (49). We used SPC solvation model and cubic boundary conditions with 12 Å buffer box size. The final system was neutral and had 0.15 M sodium chloride. A multi-stage MD simulation protocol was used, which was previously described (50). All the apo simulations were performed for a total of 1 μs. The DNA-bound structures were simulated as triplets to ensure reproducibility for 100 ns each.

### Analysis of molecular dynamics simulations

The RMSD and RMSF of protein and DNA molecule as well as the protein-ligand contacts diagram were calculated using Simulation Interactions Diagram from Schrodinger. Hydrogen bonds occupancies over the trajectories were calculated using in-house modified Schrodinger trajectory analysis python scripts. Hydrogen bonds were determined for pairs of eligible donor/acceptor atoms using criteria set by Schrodinger. For a pair of heavy atoms to form a hydrogen bond, the distance between donor-hydrogen and acceptor had to be less than 2.8 Angstrom, the angle between donor, hydrogen and acceptor had to be at most 120 degrees and the angle between hydrogen, acceptor and the next atom had to be at least 90 degrees. The residue vdW potential between A3B and DNA during the MD simulations was extracted from the simulation energies using Desmond. For both hydrogen bonds and vdW potential, errors were calculated using block averaging (51). The distance histograms display the distance between the CZ atom of Arg211 (28 in A3A) and the benzene ring centre of the side chain of Tyr315 (132 in A3A). CG, CA, CB and C of Tyr315 or Phe315 were used to determine the side chain dihedral angle. The time series of side chain conformations were generated with program VMD using 50 frames as time step (total 2000 frames).

### Cloning and mutagenesis of inactive A3B constructs

Human A3B E255A gene was codon-optimized and synthesized by GenScript. This gene was then cloned into pGEX-6p-1 vector using BamHI and EcoRI restriction sites. The pGEX-6p-1 A3B E255A catalytically inactive over-expression construct was used for all experiments in this study. All the mutations were introduced using the Q5 site-directed mutagenesis kit (NEB), and the plasmids were sequenced to verify the mutation by Genewiz.

### Protein expressions and purification

The pGEX-6p-1 A3B inactive mutant constructs were transformed into BL21 DE3 STAR E. coli strain for overexpression. Expression of GST-tagged A3B-CTD recombinant protein was performed at 17 °C for 22 hours in LB medium containing 0.5 mM IPTG and 100 μg/mL ampicillin. Cells were then pelleted, resuspended in purification buffer (50 mM Tris HCl pH 7.4; 250 mM NaCl; 0.01% Tween 20 and 1 mM DTT) and lysed with the cell disruptor. The lysate was collected and the recombinant protein was separated using a GST column. The GST tag was cleaved by the PreSecission Protease on the column at room temperature overnight. The flowthrough was then collected and further purified through size-exclusion chromatography using a HiLoad 16/60 Superdex 75 column (GE Healthcare).

### Fluorescence anisotropy-based DNA binding assay

Fluorescence anisotropy-based DNA binding assays were performed as described (28) with minor modifications. We used 5’-TAMRA labelled oligonucleotides as the binding substrate. The linear oligonuclotide sequences used were 5’-AAA-AAA-AAA-AAA-AAA-3’ (polyA) and 5’-AAA-AAA-AAT-CGA-AAA-3’ (polyA TCG). The hairpin sequences used were 5’-GCC-ATC-ATT-CGA-TGG-G-3’ (DNA hairpin) and 5’-rGrCrC-rArUrC-rUrArU-rCrGrA-rUrGrG-3’ (RNA hairpin). The reaction buffer was 50 mM Tris buffer (pH 7.4), 100 mM NaCl, 0.5 mM TCEP. The concentration of APOBEC3 was varied from 0 to 20 μM in triplicate wells containing constant amount (10 nM) of substrate. Plates were incubated for an hour on ice before reading the plates. For all experiments, fluorescence anisotropy was measured using an EnVision plate reader (PerkinElmer), with excitation at 531 nm and detecting polarized emission at 579 nm wavelength.

Data analysis was performed using Prism 7 with least-square fitting of the measured fluorescence anisotropy values (Y) at different protein concentrations (X) with a single-site binding curve with Hill slope and constant background using the equation Y=(Bmax 3 X^h)/ (Kd^h + X^h) + Background, where Kd is the equilibrium dissociation constant, h is the Hill coefficient, and Bmax is the extrapolated maximum anisotropy at complete binding. The standard deviation was calculated for each measurement point from the three independent repeats.

## Results

Despite overall similarities, the length and sequence differences of loop 1 between A3B-CTD and A3A are responsible for alterations in the active site and likely the differences in activity. The crystal structure of apo A3B-CTD reveals the active site to be in a closed conformation (42,52) compared to A3A (Figure 2A). This conformation, which results from the stacking interactions between Arg211 of loop 1 and Tyr315 of loop 7, is not compatible for DNA binding. In A3A, the side chain of Arg28 in loop 1 points out of the active site and thus results in an open conformation, in contrast to the closed conformation of A3B. The gatekeeper residue is also altered from a histidine in A3A to one of the three potential arginines in A3B for coordinating DNA binding. To elucidate the mechanisms by which DNA binding occurs, we performed multiple molecular dynamics (MD) simulations of both apo and DNA-bound structures of wildtype and variants of A3B-CTD, and A3A as a reference (**Supplementary Table S1**). These mechanisms were further validated by a series of experimental fluorescence anisotropy-based DNA binding assays of A3B variants (Table 1).

**Figure 2.**
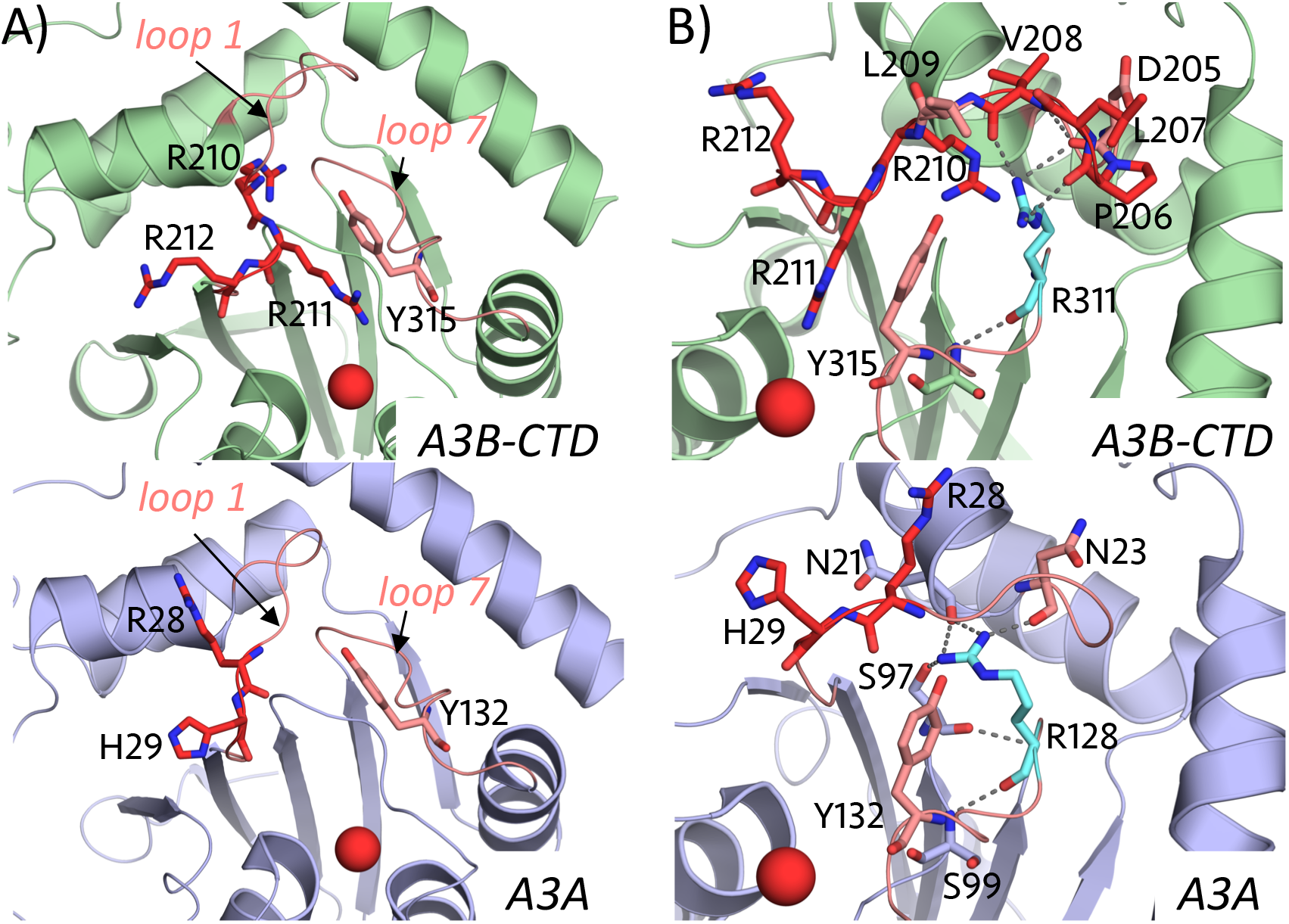
Active site of A3B-CTD versus A3A. A) A3B has a closed active site conformation while A3A has an open active site in crystal structures. B) Extra PLV residues alter the conformation of the conserved Arg311 in A3B through extensive hydrogen bond interactions. A3B-CTD (pdb: 5CQH) and A3A (pdb: 4XXO) are shown in cartoon representation (A3B in green; A3A in slate blue). The catalytic zinc is shown as red sphere. Loop 1 and loop7 are coloured salmon. 210RRR212 and 206PLV208 in A3B and 28RH29 in A3A are coloured red. The conserved arginine is coloured cyan. All the labelled residues are shown in stick representation.

**Table 1.** DNA binding affinity of A3B variants. The binding affinities, represented as K_d_, of linear DNA and hairpin DNA with TCG motif in the loop to A3B and variants, determined by fluorescence anisotropy-based assays. All A3B variants contain the E255A mutation to catalytically inactivate the enzyme and prevent substrate deamination. NB stands for no detectable binding over the range of concentrations tested (up to 20 μM of A3B); WB stands for weak binding with a K_d_ >10 μM.

| A3B variant | poly A_TCG<br>($\mu\text{M}$) | DNA hairpin<br>($\mu\text{M}$) |
| --- | --- | --- |
| WT (E255A) | $5.4 \pm 2.6$ | $2.0 \pm 0.5$ |
| P206G | NB | NB |
| Y315F | $7.3 \pm 4.6$ | $1.8 \pm 0.4$ |
| R210A | WB | WB |
| R210K | WB | WB |
| R211A | NB | NB |
| R211H | NB | NB |
| R212A | NB | $1.4 \pm 0.4$ |
| R212H | NB | $1.24 \pm 0.3$ |

### Structural mechanism of auto-inhibited conformation of apo A3B

The extra residues _206_PLV_208_ in loop 1 of A3B form a unique hydrogen bond network with Arg311 and Asp205 (Figure 2B). Specifically, the backbone carbonyl oxygen of Pro206 makes a hydrogen bond with NH2 atom of Arg311; Val208 forms two backbone hydrogen bonds with the backbone of Asp205 and NH1 atom of Arg311. The backbone carbonyl oxygen of Asp205 also has a hydrogen bond with NH1 atom of Arg311. Arg311 is conserved among all A3 domains (**Supplementary Figure S1A**); however when we superimposed all the available active apo A3 structures (A3A, A3B-CTD, A3C, A3F-CTD and A3G-CTD), the sidechain of this conserved Arg was locked in a hydrogen bond network distinct from that in A3B. (Figure 2B and **Supplementary Figure S1B**). This network involves primarily the backbone atoms of conserved Ser97, Ser99, Asn21 (Gln in A3C and His in A3F-CTD and A3G-CTD) and Asn23 (Lys in A3C, A3F-CTD and A3G-CTD). In A3A, the side chain of Ser97 forms an additional hydrogen bond with the conserved Arg. Instead in A3B, these residues form a similar hydrogen bond network with Arg210 in loop 1. The distinct conformation of Arg311 in A3B-CTD and hydrogen bonding with the _206_PLV_208_ in loop 1 might contribute to the closed conformation of A3B-CTD as well as A3B’s lower activity. To investigate _206_PLV_208_’s role in regulating activity, 1 μs MD simulations were performed on A3A and A3B-CTD as well a variant where the _206_PLV_208_ sequence was deleted (A3B-CTD-ΔPLV). As neither the crystal structure of apo A3A nor A3B-CTD were completely of the wild type sequences (the apo A3A structure has an inactive mutation while apo A3B-CTD has a truncated loop 3 and four additional mutations to increase solubility), the wild type A3A and A3B-CTD structures were modelled.

From the MD simulation trajectories, we first examined the stability of the closed active site conformation (Figure 3). In wild type A3B, the distance between the side chain of Arg211 in loop 1 and Tyr315 in loop 7 varied around 6 Å, which is within the range of stacking interactions that close the active site. The closed active site conformation in A3B was stable during the MD simulations, with the distance not increasing above 10 Å. In contrast, the equivalent residues in wild type A3A, Arg28 and Tyr132 had a distance distribution around 15 Å, which indicates the active site is in the open conformation (Figure 3A and C). Interestingly, in A3B-CTD-ΔPLV, Arg211 lost the stacking interactions with Tyr315. The distance between the side chains of Arg211 and Tyr315 was more than 12 Å during the MD simulations. As a result, the active site conformation was altered into the open conformation, analogous to that observed in A3A. The more open active site correlates with higher activity in A3A and A3B-CTD-ΔPLV compared to wild type A3B.

**Figure 3.**
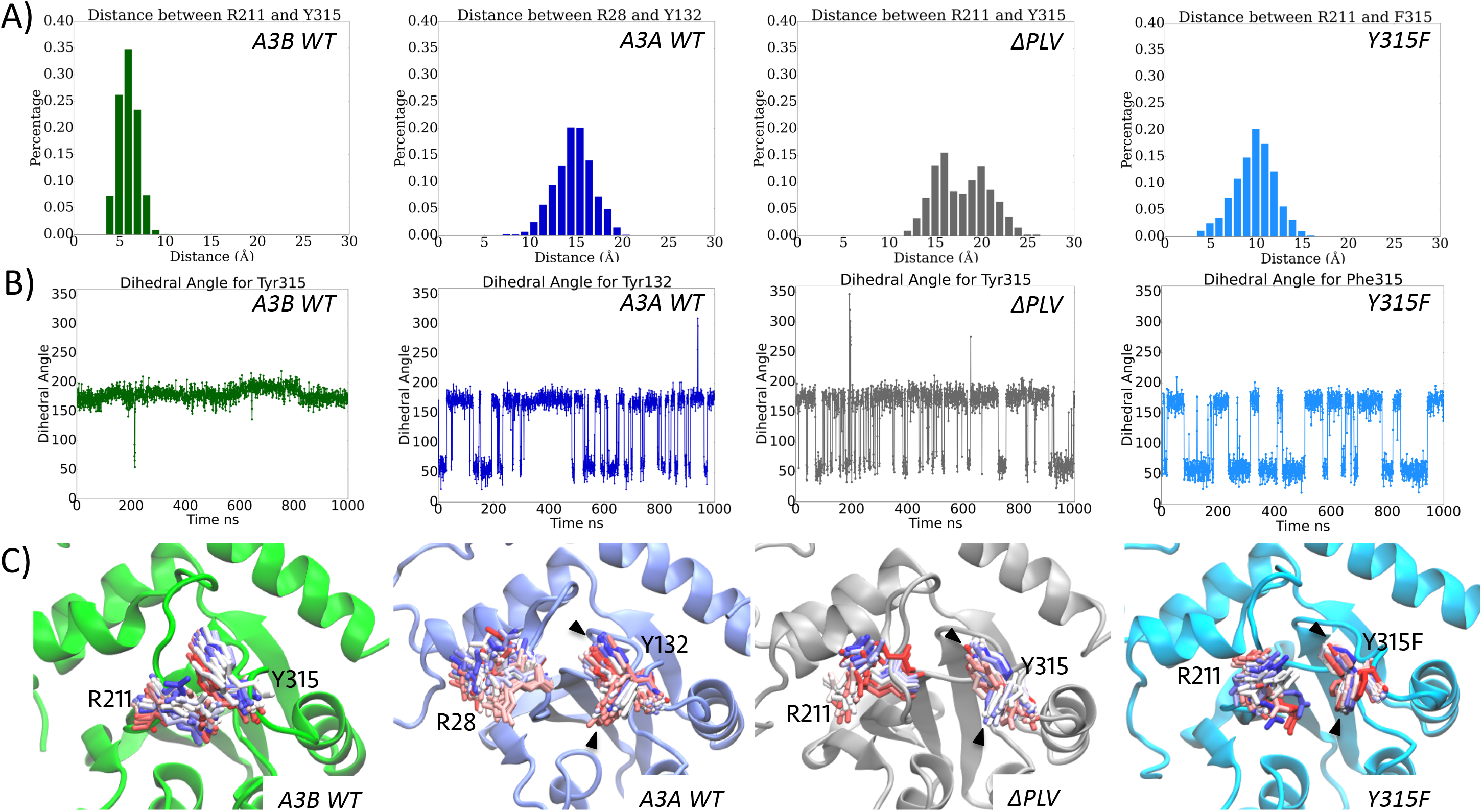
Dynamics of the active site in A3A, A3B and A3B mutants. A) The histogram of the distance between CZ atom of Arg311 (Arg28 in A3A) and centre of benzene ring of Tyr315 (Tyr132 in A3A) in wild type A3B, A3A, A3B ΔPLV and Y315F mutants. B) The dihedral angle of the side chain (C, CA, CB and CG) of Tyr315 (Tyr132 in A3A) over 1 μs MD simulations in wild type A3B, A3A, A3B ΔPLV and Y315F mutants. C) The time series of the side chain conformations of Arg311 (Arg28 in A3A) and Tyr315 (Tyr132 in A3A) (shown as stick; coloured based on the simulation time, start as red and end with blue) during 1 μs MD of wild type A3B (green), A3A (slate blue), A3B ΔPLV (grey) and Y315F (cyan) variants. Different conformations of Tyr315 (Y315F) are indicated with arrows.

We also monitored the side chain conformation of Tyr315, which is analogous to Tyr132 in A3A, during the MD simulations as an indicator for the compatibility to bind DNA (Figure 3B and C). The Tyr side chain has to undergo a conformer change to accommodate binding of DNA at the active site. In wild type A3B, the side chain dihedral angle χ^1^ of Tyr315 remained as ~ 180° in 99.8% of the MD simulations time, in agreement with our finding that the closed active site conformation in A3B is not compatible for DNA binding. The same dihedral angle χ^1^ of Tyr132 in A3A changed from about 180° to 60° upon DNA binding in the crystal structure (46). In the MD simulation, the side chain of Tyr132 in A3A sampled between the apo and DNA bound conformations (68.0% of the time in DNA-bound conformation). In A3B-CTD-ΔPLV, the side chain of Tyr315 sampled two conformations (72.8% of time in DNA-bound conformation) as in A3A. The high sampling frequency of DNA-compatible chain conformation of Tyr315 in A3B-CTD-ΔPLV and A3A may also account for the higher activity compared to wild type A3B. Throughout the MD trajectory of wild type A3B, the _206_PLV_208_ hydrogen bond network interacted with Tyr315, retaining the auto-inhibited conformation (**Supplementary Figure S2A**). Specifically, the OH atom of Tyr315 interacted with both NH2 atom of Arg311 and backbone carbonyl oxygen of Pro206 through direct and water-mediated hydrogen bonds. As a result, the side chain of Tyr315 was locked in the DNA incompatible conformation (Figure 3B and C). These hydrogen bonds were stable throughout the MD simulations (**Supplementary Figure S2B**). To verify the role of this auto-inhibited conformation in down-regulating A3B’s activity, the Y315F variant was modelled and subjected to the same 1 μs MD simulations. Phe315 in the Y315F variant lost the ability to interact with _206_PLV_208_ hydrogen bond network, as the hydroxyl group was lost, and was released from the auto-inhibited conformation. The side chain dihedral angle χ^1^ of Phe315 sampled the DNA compatible conformation with 49.2% frequency in MD simulations (Figure 3B). However, the stacking interaction between Arg211 and Phe315 was maintained during the MD simulations and thus the Y315F variant still kept the active site in the closed conformation (Figure 3A and C). Hence, disrupting the hydrogen bonding network between residue 315 and _206_PLV_208_ shifted this residue to a DNA-compatible conformer but was not enough to open up the active site.

Both A3B-CTD and A3G-CTD have longer loop 1, which includes a proline residue, compared to A3A. Considering its unique geometry and rigidity compared to other amino acids, this extra proline may help stabilize the conformation of a longer loop 1. To test this hypothesis, we modelled A3B P206G mutant and examined the root-mean-squared-fluctuations (RMSFs) of loop 1 as a measure of dynamics (Figure 4). Despite the differences in activity, wild type A3B and A3A have similar levels of loop 1 dynamics with RMSF values varying around 4 Å, as well as wild type A3G-CTD (**Supplementary Figure S3**). The RMSFs of loop 1 in A3B Y315F mutant and A3B-CTD-ΔPLV were also within 4 Å during the MD simulations, which suggests the importance and consistency of loop 1 dynamics in DNA binding. Loop 1 in A3B P206G variant, however, is highly dynamic with RMSFs varying from 1 up to 8 Å indicating that loop 1 bearing the P2016G mutation may not be able to stably coordinate DNA.

**Figure 4.**
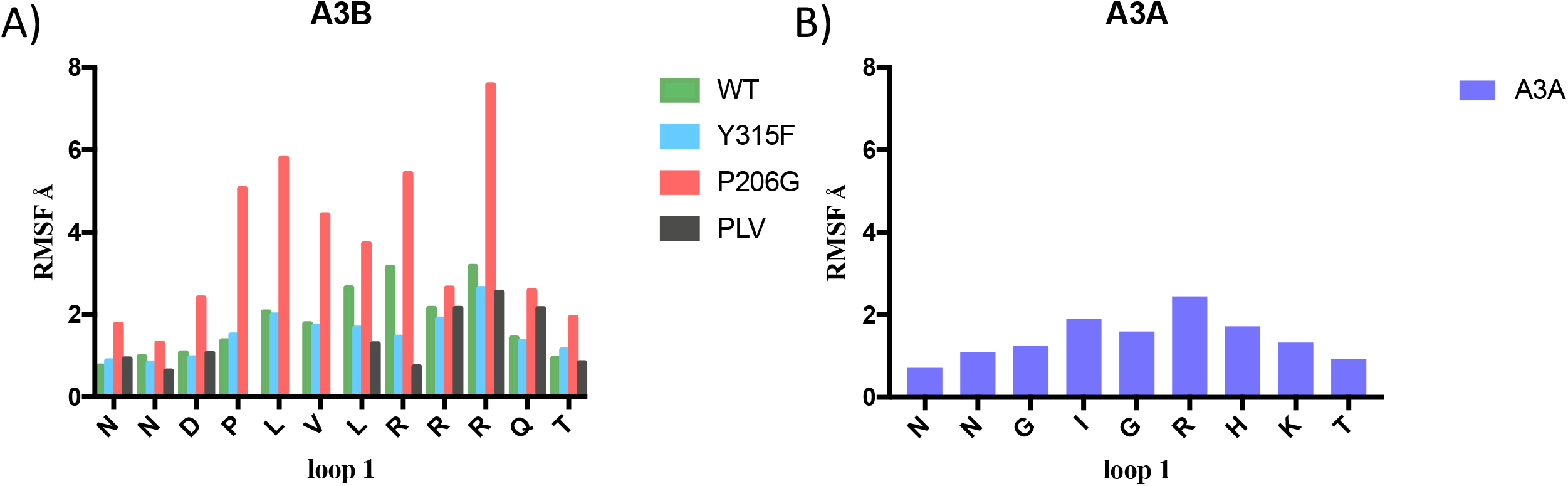
The fluctuations of loop 1 during MD simulations. A) The root-mean-squared-fluctuations (RMSFs) of individual residues of loop 1 in wild type A3B and A3B variants. B) The RMSF of all residues in loop 1 of wild type A3A.

To validate our computational hypotheses, mutations were introduced into A3B and experimental fluorescence anisotropy-based DNA binding assays were performed (Table 1). The low binding affinity and catalytic activity of wild type A3B poses challenges in assessing changes in deamination rates and DNA binding. Nevertheless, we were able to show that the A3B inactive variant can bind to both linear (Kd ~ 5.4 μM) and hairpin DNA that has substrate sequence TCG in the stem loop (Kd ~2.0 μM). Y315F variant bound to DNA with similar levels as wild type, which is in agreement with our computational findings that Y315F maintained the closed active site conformation as in wild type. In contrast, the P206G variant lost the ability to bind both linear and hairpin DNA. Recent studies have shown that removing PLV from loop 1 in A3B increases the enzyme’s activity, (48) which is in agreement with the more open active site conformation that we observed for A3B-CTD-ΔPLV. Thus, the closed active site conformation observed in modelling and simulations were in complete agreement with experimental binding and catalytic activity. Together these findings strongly suggest that the extra PLV residues in loop 1 is the key for restricting and regulating A3B’s deamination activity, with the proline stabilizing the conformation of the longer loop 1 for DNA binding.

### Molecular mechanism of A3B-DNA recognition

The role of loop 1 and molecular mechanism of DNA binding had remained unknown for A3B, as recently determined A3B-CTD DNA co-crystal structure is a chimera with the crucial loop 1 swapped from A3A. Here we resorted to careful molecular modelling using available crystal structures to answer for A3B: 1. How does DNA bind to A3B? 2. Which residue is the gatekeeper for latching DNA in the active site? 3. How does A3B define its substrate specificity for thymidine over cytidine at -1 position? To address these questions, we modelled fully wild type A3B-CTD bound to substrate DNA containing a TCG trinucleotide motif based on the crystal structures of apo A3B-CTD and A3A-DNA complex (see Methods for details). The quality of the models was further examined through both computational analysis of triplicate 100 ns MD simulations of A3A and A3B complexes (**Supplementary Table S1**), and experimental DNA binding assays of inactive A3B variants (Table 1). All the MD simulations converged and were stable over the 100 ns trajectory time.

### Arg211 is the gatekeeper residue sequestering DNA in the active site

As A3B-CTD has a unique _210_RRR_212_ patch in loop 1 instead of _28_RH_29_ compared to A3A, and either Arg211 or Arg212 could be the gatekeeper for DNA binding. To identify the critical residue, we generated DNA-bound models with either Arg211 or Arg212 latching DNA in the active site and performed and analysed triplicate 100 ns MD simulations. The Arg212 model DNA complex was much less stable during the MD simulations compared to either A3A or Arg211 model as indicated by the considerable larger RMSFs especially at the two termini of bound DNA (Figure 5A). In both A3A structure and Arg211 model, Glu255 consistently interacted with substrate cytidine 98.94% and 96.29% of the simulation time, respectively (Figure 5B). However in the Arg212 model, the contact frequency was decreased to only 60.21%, which suggests poor quality of the model. Thus the stability over MD simulations indicated that Arg211 rather than Arg212 is the gatekeeper for DNA binding in A3B.

**Figure 5.**
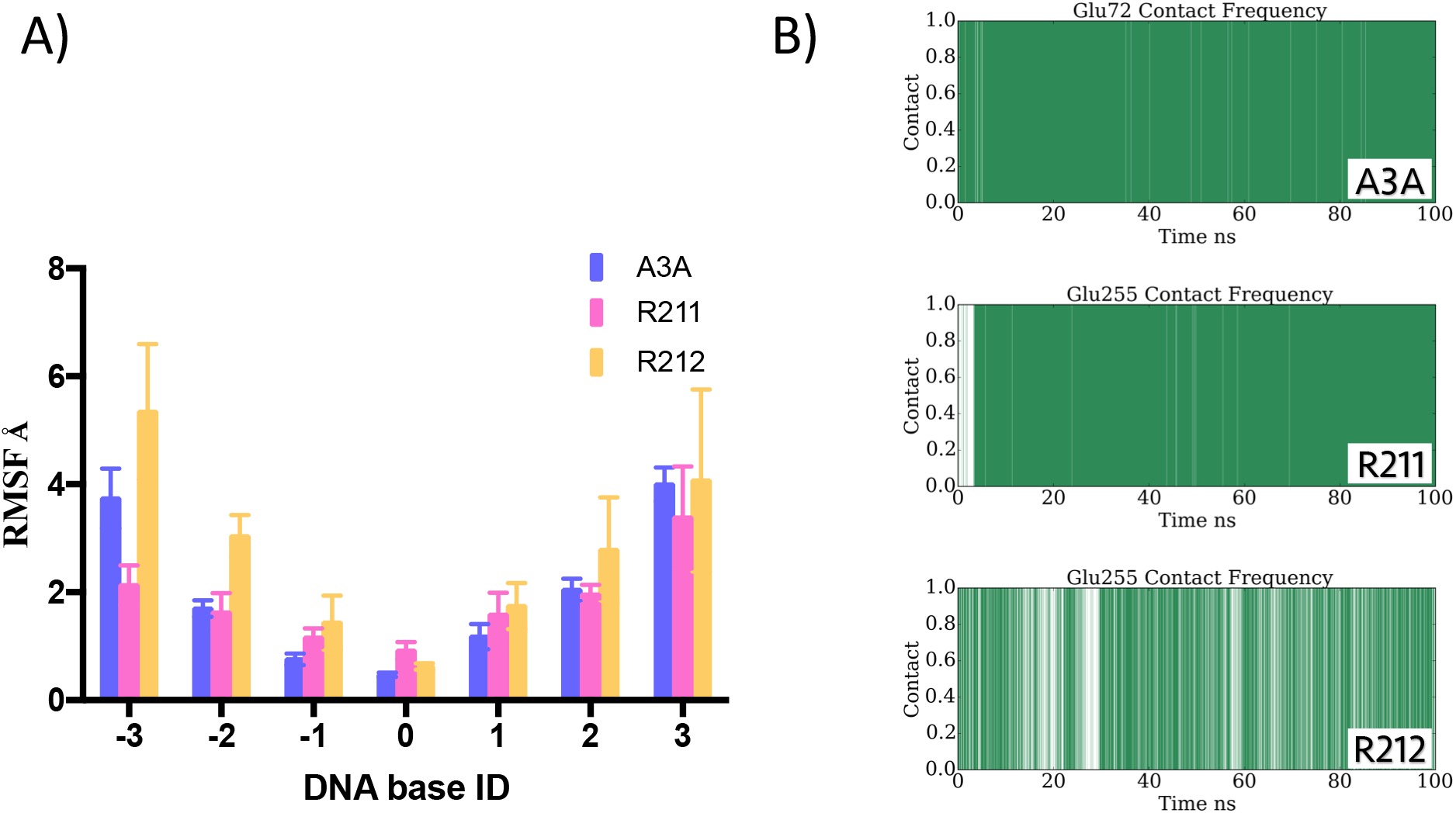
Comparison of A3B-DNA model structures with either R211 or R212 latching the DNA in the active site. A) The root-mean-squared-fluctuations (RMSFs) of individual bases of DNA molecule in A3A, A3B R211 and R212 models. B) The contact frequency of the intermolecular interactions between catalytic residue Glu255 and substrate cytidine base over time in A3A, A3B R211 and R212 models.

To further verify our results from computational analysis, we experimentally generated A3B R210A, R210K, R211A, R211H, R212A and R212H inactive mutants and tested DNA binding using fluorescence anisotropy-based binding assay (Table 1). Both R210A and R210K mutants were able to bind DNA but with decreased binding affinity, which is in agreement with Arg210’s role in stabilizing overall structure through the conserved hydrogen bond network. R212A and R212H variants bound to DNA with same affinity as wild type A3B, which confirmed that Arg212 is not the gatekeeper for DNA binding. In contrast, R211A and R212A mutants lost binding completely for both ssDNA and hairpin DNA. Recent A3B activity studies have also shown that R211A mutant lost deamination activity (42). Thus experimental binding assay data were in agreement with our model and finding that Arg211 is the critical residue for DNA binding.

### ssDNA binding to A3B-CTD

The DNA-bound model of A3B-CTD revealed the molecular mechanism as well as the role of loop 1 for DNA binding to A3B. Overall, ssDNA bound to A3B-CTD in a U-shape similar to A3A (Figure 6A). Loop 1 underwent major conformational changes to open up the active site for DNA binding (Figure 6B), especially at Arg211 which stacks against Tyr315 in the closed active site conformation. In addition, the side chain of Tyr315 rotated from ~180° to ~60° as in A3A to accommodate DNA binding (Figure 6B). In general, our model overlaid well with A3B chimera DNA co-crystal structure, but also provided additional information for loop 1 in DNA binding (Figure 6C). The loops around the active site, loop 1, 3 and 7, have extensive vdW interactions with ssDNA (Figure 6D and E), especially loop 1 which contributed up to 50% of the total vdW contacts. The most critical intermolecular interaction between A3B and ssDNA involved the gatekeeper Arg211. Arg211 coordinated DNA binding through both hydrogen bond interactions with the phosphate backbone of -1 T, 0 C and +1 G bases and hydrophobic interactions with DNA backbone (Figure 7B). Arg212 instead stabilized DNA binding through either stacking (Figure 7A) or hydrogen bond interactions with +1 G base.

**Figure 6.**
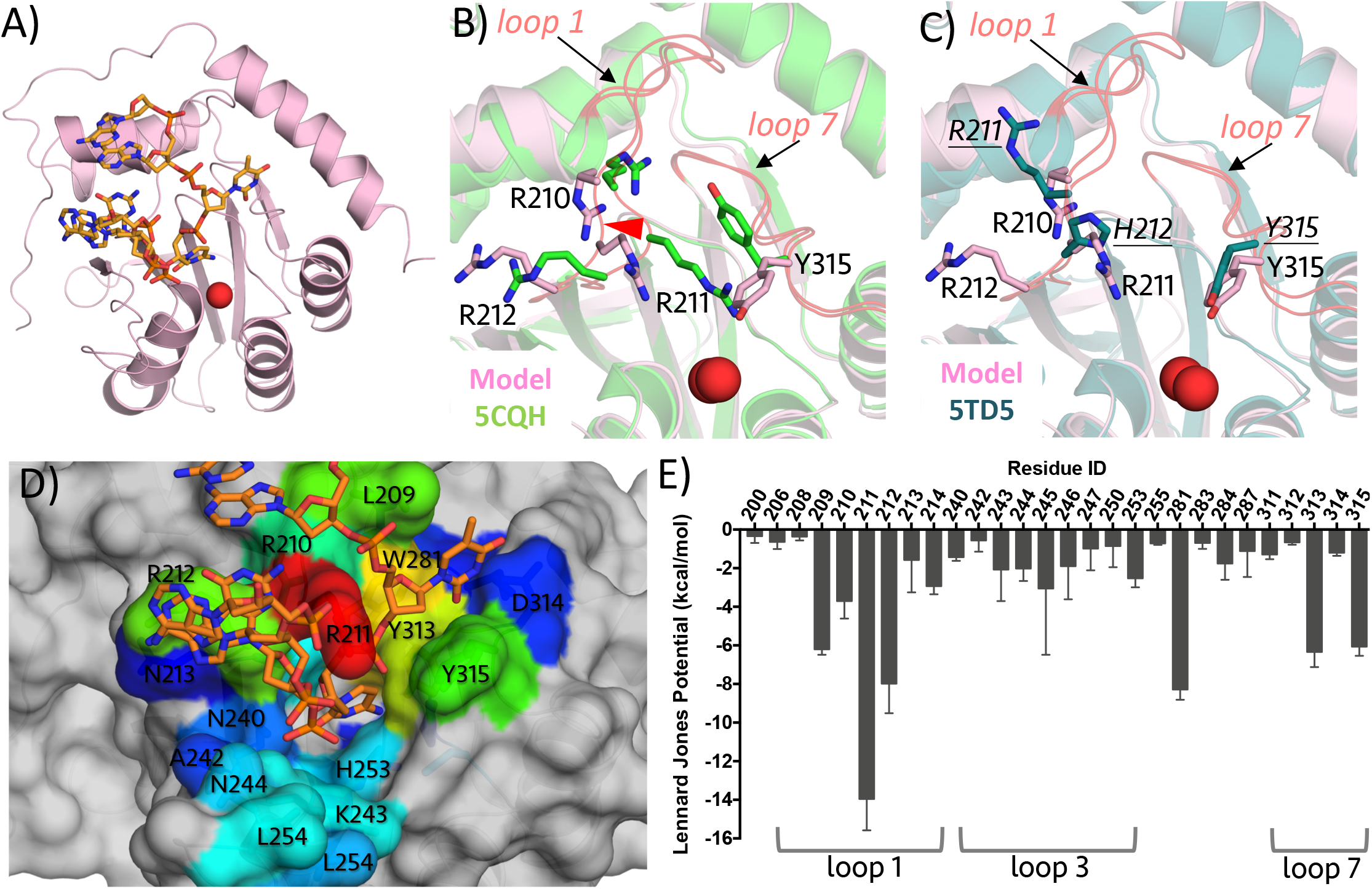
Structural model of A3B-CTD in complex with ssDNA. A) Overall structure for A3B-CTD DNA model. B) Conformational changes of residues R210, R211, R212 and Y315 upon DNA binding, with side chains displayed in stick representation. Loop 1 and loop7 are coloured salmon and the conformational changes upon DNA binding are indicated by arrows. C) Structural comparison between our model and A3B chimera DNA crystal structure (PDB: 5TD5). D,E) Mean vdW contacts between protein and ssDNA over triplicate MD trajectories. The residues are coloured on a rainbow scale from blue to red for increasing contacts; hence warmer colours indicate residues with the most contribution to the intermolecular contacts. The cut-off for the scale is -0.5 kcal/mol.

### D314 defines substrate specificity for thymidine over cytidine at -1 position

In A3A, the substrate specificity for thymidine over cytidine at -1 position is determined through hydrogen bond interactions with Asp132 (46). In our A3B-DNA model, we observed the same hydrogen bonding pattern between -1 T base and Asp314 as in A3A (Figure 7D and **Supplementary Figure S5A**). Specifically, O2 atom of -1 T formed a direct hydrogen bond with Asp314 backbone, while OD1 and OD2 atoms of Asp314 had both direct and water-mediated hydrogen bonding with N3 and O4 of -1 T base. All these hydrogen bond interactions were stable during MD simulations (**Supplementary Figure S5B**). In contrast, when thymidine was changed to a cytidine, the side chain hydrogen bond interactions between Asp314 and -1 T were disrupted. As a result, DNA containing -1 C was more dynamic in the active site and sampled alternate conformations (**Supplementary Figure S5C and D**). Similarly, in the Arg212 model, which we deduced to be poor based on dynamics above, Asp314 had diverse and unstable hydrogen bond interactions with -1 T (**Supplementary Figure S4A and B**). These findings suggest that A3B likely uses the same molecular mechanism to determine the substrate specificity as A3A at -1 position since Asp314 is conserved and the DNA interactions maintained between the two A3s.

**Figure 7.**
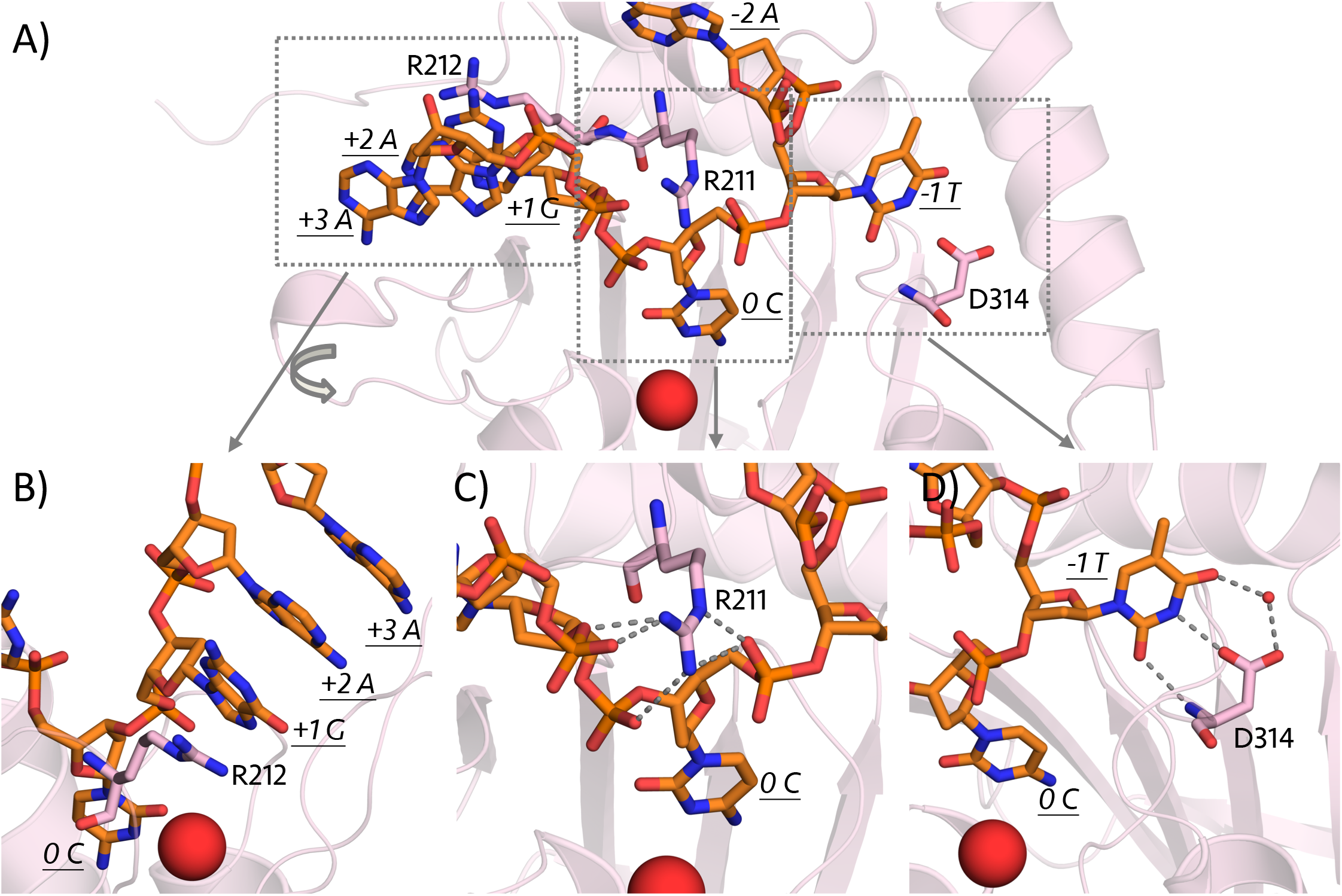
Intermolecular interactions between A3B and ssDNA. A) Overview of key residues in binding to DNA. B) R212 can stack with downstream DNA bases. C) R211 forms extensive hydrogen bond interactions with DNA backbone. D) D314 makes extensive hydrogen bonds with -1 base that defines substrate specificity. The final frame of the MD simulation is displayed. The protein is displayed as a pink-coloured ribbon diagram and the bound DNA is in orange stick representation. The zinc ion at the active site is depicted as a red-coloured sphere. The side chains of R210, R211, R212, D314 and Y315 are shown as sticks. The hydrogen bond interactions are indicated with grey dashed lines.

## DISCUSSION

A3 proteins have the same overall fold, with highly conserved active sites and yet neither the available structures nor the amino acid sequences offer obvious insights into why they have highly varying catalytic activity, from totally inactive pseudo-catalytic NTDs to the highly active A3A. Instead the seemingly minor diversity in the loops 1, 3 and 7 around the active site are responsible for regulating A3 activity and function in innate immunity and cancer development. This study identified the molecular mechanism by which A3B structurally downregulates deamination activity compared to other A3s, especially the highly homologous A3A.

In this study, we revealed the critical role of loop 1 in regulating A3B activity and DNA binding. The _206_PLV_208_ insertion in loop 1 causes a closed active site conformation, which blocks DNA binding and thus substrate deamination. Our results suggest that the length and sequence differences in loop 1, which were missing in the DNA-bound A3B crystal structure (47), are key in regulating activity of Z1 domain A3s (Figure 8). A short loop 1, as in A3A or A3B-CTD-ΔPLV, results in high catalytic activity. A longer loop 1 with proline, which stabilizes the overall loop 1 conformation in A3G-CTD, results in medium activity. Finally, having the auto-inhibited conformation and a longer loop 1 which forms molecular interactions with loop 7 in A3B-CTD further restricts deamination activity. In contrast to all the other A3s, in A3B the conserved Arg311 participates in a distinct hydrogen bond network involving the PLV insertion, and Arg211 stacks against Tyr315 to further restrict A3B’s activity. Recent work also revealed the role of loop 7 in defining substrate specificity, as swapping loop 7 of A3A into A3G reversed the substrate specificity (53). Thus, the detailed analysis of A3B structure revealed insights into how amino acid changes around the active site can structurally regulate the relative catalytic activity of A3s despite highly similar overall structure and conserved active site.

**Figure 8.**
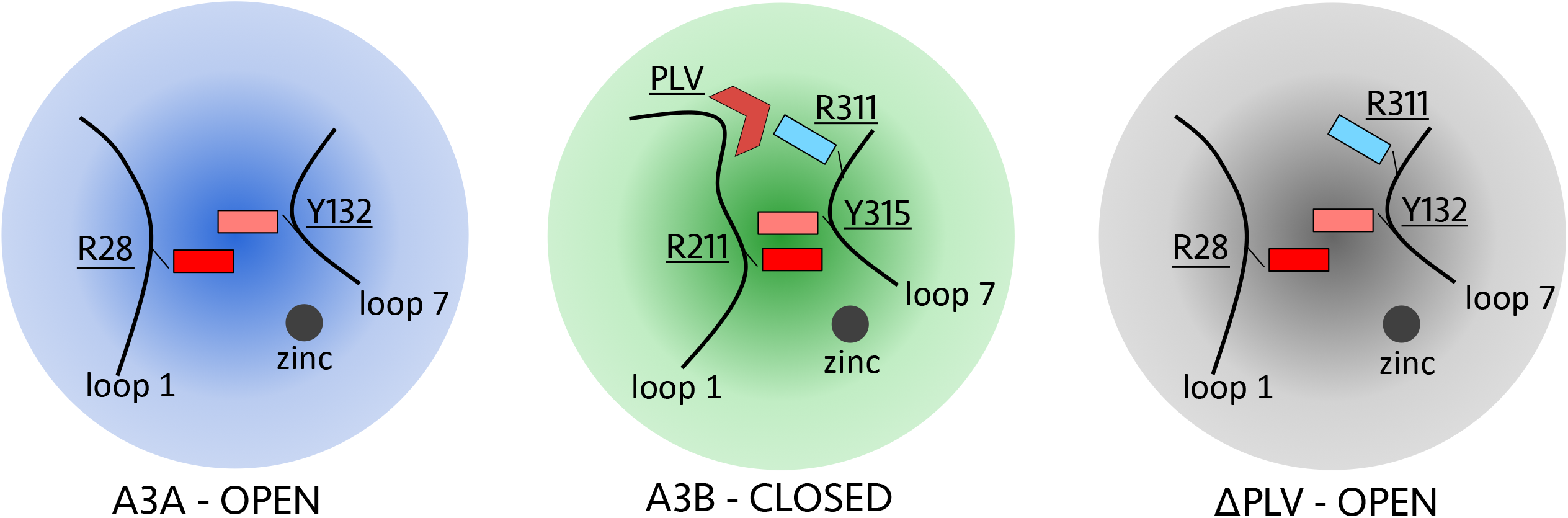
A schematic representation of the mechanism by which A3B regulates activity. The structural features at the active site of A3A, A3B-CTD and PLV deletion variant that regulate catalytic activity. Loop 1 and loop 7 are shown as lines, and the catalytic zinc is represented as a grey sphere. The side chain of R28, Y132 in A3A and R211, Y315, R311 in A3B are shown as rectangles. PLV in A3B is represented as a wedge.

Our structural model for DNA-bound wild type A3B, which was experimentally validated, further reveals the role of loop 1 in DNA binding. Our model structure provides a comprehensive mechanism for DNA recognition, the key residue for substrate specificity at -1 position, and the crucial gatekeeper Arg211 which coordinates DNA in the active site. In A3A, His29 is the gatekeeper for DNA binding through both hydrogen bond interactions to DNA backbone and stacking interactions to +1 base. A3B R211H variant, however, showed no binding to DNA (Table 1). Rather the two types of interactions His29 makes with bound DNA in A3A can be assigned separately to Arg211 and Arg212 in A3B, as Arg211 hydrogen bonds to DNA in the active site while Arg212 stabilizes DNA binding through either stacking or hydrogen bond interactions with the +1 DNA base. Experimentally, A3B R212H variant was still able to bind DNA with the same affinity (Kd ~ 1.34 μM) as wild type (Table 1), which is in agreement with this mode of binding. A3B should be able to bind pre-bent or hairpin DNA like A3A (BioRxiv: https://doi.org/10.1101/176297) based on our model, and likely with higher affinity as the entropic cost of bending the DNA would be decreased. Accordingly, we observed higher binding affinity to hairpin DNA (2.0 μM) compared to linear DNA (5.4 μM) for A3B (Table 1). Interestingly, unlike A3A (BioRxiv: https://doi.org/10.1101/176297) (54), A3B showed no binding to RNA hairpin. These structural and functional differences between A3A and A3B despite their high sequence similarity might have implications for their biological function as well as cellular localization.

There are still several A3 domains with structures not determined, let alone two-domain constructs of A3s. The low solubility and DNA affinity of certain A3 proteins have required introducing mutations to be able to structurally and biochemically characterize these proteins in vitro (25-27,29-31,38,42,45-47), or altogether prevented such characterization especially for NTDs. For instance, wild type A3B-CTD has poor solubility and low binding affinity towards DNA, which makes crystalizing native A3B-DNA complex extremely challenging. The available DNA-bound structure was that of an A3B chimera engineered to increase affinity and promote crystallization. While this structure did not inform on the role of loop 1 in DNA binding, the apo structure of A3B with the native loop1 and A3A-DNA structures enabled computational modelling of wild type A3B bound to substrate DNA. Future studies focusing on the active site loops around A3 family members will elucidate the differences between the catalytically active and pseudo-catalytic A3 domains. Similarly, combining experimental structures with computational modelling, verified by simulations and experimental mutational analysis, can provide insights into the function and DNA binding of other A3s

Designing inhibitors or activators for A3s has proven to be extremely challenging. Our results provide opportunities for drug design to specifically target A3B and thus benefit cancer therapeutics. Small molecules that stabilize the unique auto-inhibited mode of A3B might be able to allosterically inhibit A3B without cross-reacting with other A3s. Besides, the residue-specific information on regulation of auto-inhibition and closed active site conformation provides the starting point for engineering A3s domains to achieve varying catalytic efficiencies or distinct substrate specificity.

## Acknowledgement

The authors would like to thank Dr. Mohan Somasundaran for helpful advice and suggestions.

## References

1. Betts, L., Xiang, S., Short, S.A., Wolfenden, R. and Carter, C.W., Jr. (1994) Cytidine deaminase. The 2.3 A crystal structure of an enzyme: transition-state analog complex. j. Mol. Biol., 235, 635–656.

2. Jarmuz, A., Chester, A., Bayliss, J., Gisbourne, J., Dunham, I., Scott, J. and Navaratnam, N. (2002) An anthropoid-specific locus of orphan C to U RNA-editing enzymes on chromosome 22. Genomics, 79, 285–296.

3. Wedekind, J.E., Dance, G.S., Sowden, M.P. and Smith, H.C. (2003) Messenger RNA editing in mammals: new members of the APOBEC family seeking roles in the family business. Trends Genet., 19, 207–216.

4. Conticello, S.G., Thomas, C.J., Petersen-Mahrt, S.K. and Neuberger, M.S. (2005) Evolution of the AID/APOBEC family of polynucleotide (deoxy)cytidine deaminases. Mol. Biol. Evol., 22, 367–377.

5. LaRue, R.S., Andresdottir, V., Blanchard, Y., Conticello, S.G., Derse, D., Emerman, M., Greene, W.C., Jonsson, S.R., Landau, N.R., Lochelt, M. et al. (2009) Guidelines for naming nonprimate APOBEC3 genes and proteins. J. Virol., 83, 494–497.

6. Sheehy, A.M., Gaddis, N.C., Choi, J.D. and Malim, M.H. (2002) Isolation of a human gene that inhibits HIV-1 infection and is suppressed by the viral Vif protein. Nature, 418, 646–650.

7. Zheng, Y.H., Irwin, D., Kurosu, T., Tokunaga, K., Sata, T. and Peterlin, B.M. (2004) Human APOBEC3F is another host factor that blocks human immunodeficiency virus type 1 replication. J. Virol., 78, 6073–6076.

8. Dang, Y., Siew, L.M., Wang, X., Han, Y., Lampen, R. and Zheng, Y.H. (2008) Human cytidine deaminase APOBEC3H restricts HIV-1 replication. j. Biol. Chem., 283, 11606–11614.

9. Dang, Y., Wang, X., Esselman, W.J. and Zheng, Y.H. (2006) Identification of APOBEC3DE as another antiretroviral factor from the human APOBEC family. j. Virol., 80, 10522–10533.

10. Bogerd, H.P., Wiegand, H.L., Doehle, B.P., Lueders, K.K. and Cullen, B.R. (2006) APOBEC3A and APOBEC3B are potent inhibitors of LTR-retrotransposon function in human cells. Nucleic Acids Res., 34, 89–95.

11. Muckenfuss, H., Hamdorf, M., Held, U., Perkovic, M., Lower, J., Cichutek, K., Flory, E., Schumann, G. G. and Munk, C. (2006) APOBEC3 proteins inhibit human LINE-1 retrotransposition. j. Biol. Chem., 281, 22161–22172.

12. Burns, M.B., Lackey, L., Carpenter, M.A., Rathore, A., Land, A.M., Leonard, B., Refsland, E.W., Kotandeniya, D., Tretyakova, N., Nikas, J.B. et al. (2013) APOBEC3B is an enzymatic source of mutation in breast cancer. Nature, 494, 366–370.

13. Taylor, B.J., Nik-Zainal, S., Wu, Y.L., Stebbings, L.A., Raine, K., Campbell, P.J., Rada, C., Stratton, M.R. and Neuberger, M.S. (2013) DNA deaminases induce break-associated mutation showers with implication of APOBEC3B and 3A in breast cancer kataegis. Elife, 2, e00534.

14. Starrett, G.J., Luengas, E.M., McCann, J.L., Ebrahimi, D., Temiz, N.A., Love, R.P., Feng, Y., Adolph, M.B., Chelico, L., Law, E.K. et al. (2016) The DNA cytosine deaminase APOBEC3H haplotype I likely contributes to breast and lung cancer mutagenesis. Nat Commun, 7, 12918.

15. Burns, M.B., Temiz, N.A. and Harris, R.S. (2013) Evidence for APOBEC3B mutagenesis in multiple human cancers. Nat. Genet., 45, 977–983.

16. Wissing, S., Montano, M., Garcia-Perez, J.L., Moran, J.V. and Greene, W.C. (2011) Endogenous APOBEC3B restricts LINE-1 ret rot rans position in transformed cells and human embryonic stem cells. J. Biol. Chem., 286, 36427–36437.

17. Xu, R., Zhang, X., Zhang, W., Fang, Y., Zheng, S. and Yu, X.F. (2007) Association of human APOBEC3 cytidine deaminases with the generation of hepatitis virus B x antigen mutants and hepatocellular carcinoma. Hepatology, 46, 1810–1820.

18. Bonvin, M. and Greeve, J. (2007) Effects of point mutations in the cytidine deaminase domains of APOBEC3B on replication and hypermutation of hepatitis B virus in vitro. J. Gen. Virol., 88, 3270–3274.

19. Leonard, B., Hart, S.N., Burns, M.B., Carpenter, M.A., Temiz, N.A., Rathore, A., Vogel, R.I., Nikas, J.B., Law, E.K., Brown, W.L. et al. (2013) APOBEC3B Upregulation and Genomic Mutation Patterns in Serous Ovarian Carcinoma. Cancer Res., 73, 7222–7231.

20. Sieuwerts, A.M., Willis, S., Burns, M.B., Look, M.P., Meijer-Van Gelder, M.E., Schlicker, A., Heideman, M.R., Jacobs, H., Wessels, L., Leyland-Jones, B. et al. (2014) Elevated APOBEC3B correlates with poor outcomes for estrogen-receptor-positive breast cancers. Horm. Cancer, 5, 405–413.

21. Roberts, S.A., Lawrence, M.S., Klimczak, L.J., Grimm, S.A., Fargo, D., Stojanov, P., Kiezun, A., Kryukov, G.V., Carter, S.L., Saksena, G. et al. (2013) An APOBEC cytidine deaminase mutagenesis pattern is widespread in human cancers. Nat. Genet., 45, 970–976.

22. Periyasamy, M., Singh, A.K., Gemma, C., Kranjec, C., Farzan, R., Leach, D.A., Navaratnam, N., Palinkas, H.L., Vertessy, B.G., Fenton, T.R. et al.1 (2017) p53 controls expression of the DNA deaminase APOBEC3B to limit its potential mutagenic activity in cancer cells. Nucleic Acids Res., 45, 11056–11069.

23. Harris, R.S. (2015) Molecular mechanism and clinical impact of APOBEC3B-catalyzed mutagenesis in breast cancer. Breast Cancer Res., 17, 8.

24. Kidd, J.M., Newman, T.L., Tuzun, E., Kaul, R. and Eichler, E.E. (2007) Population stratification of a common APOBEC gene deletion polymorphism. PLoS Genet., 3, e63.

25. Chen, K.M., Harjes, E., Gross, P.J., Fahmy, A., Lu, Y., Shindo, K., Harris, R.S. and Matsuo, H. (2008) Structure of the DNA deaminase domain of the HIV-1 restriction factor APOBEC3G. Nature, 452, 116–119.

26. Harjes, E., Gross, P.J., Chen, K.M., Lu, Y., Shindo, K., Nowarski, R., Gross, J.D., Kotler, M., Harris, R.S. and Matsuo, H. (2009) An extended structure of the APOBEC3G catalytic domain suggests a unique holoenzyme model. J. Mol. Biol., 389, 819–832.

27. Shandilya, S.M., Nalam, M.N., Nalivaika, E.A., Gross, P.J., Valesano, J.C., Shindo, K., Li, M., Munson, M., Royer, W.E., Harjes, E. et al. (2010) Crystal structure of the APOBEC3G catalytic domain reveals potential oligomerization interfaces. Structure, 18, 28–38.

28. Li, M., Shandilya, S.M., Carpenter, M.A., Rathore, A., Brown, W.L., Perkins, A.L., Harki, D.A., Solberg, J., Hook, D.J., Pandey, K.K. et al. (2012) First-in-class small molecule inhibitors of the single-strand DNA cytosine deaminase APOBEC3G. ACS Chem. Biol., 7, 506–517.

29. Bohn, M.F., Shandilya, S.M., Albin, J.S., Kouno, T., Anderson, B.D., McDougle, R.M., Carpenter, M.A., Rathore, A., Evans, L., Davis, A.N. et al. (2013) Crystal structure of the DNA cytosine deaminase APOBEC3F: the catalytically active and HIV-1 Vif-binding domain. Structure, 21, 1042–1050.

30. Bohn, M.F., Shandilya, S.M., Silvas, T.V., Nalivaika, E.A., Kouno, T., Kelch, B.A., Ryder, S.P., Kurt-Yilmaz, N., Somasundaran, M. and Schiffer, C.A. (2015) The ssDNA Mutator APOBEC3A Is Regulated by Cooperative Dimerization. Structure, 23, 903–911.

31. Kouno, T., Luengas, E.M., Shigematsu, M., Shandilya, S.M., Zhang, J., Chen, L., Hara, M., Schiffer, C.A., Harris, R.S. and Matsuo, H. (2015) Structure of the Vif-binding domain of the antiviral enzyme APOBEC3G. Nat. Struct. Mol. Biol., 22, 485–491.

32. Chelico, L., Pham, P., Calabrese, P. and Goodman, M.F. (2006) APOBEC3G DNA deaminase acts processively 3’ —> 5’ on single-stranded DNA. Nat. Struct. Mol. Biol., 13, 392–399.

33. Holden, L.G., Prochnow, C., Chang, Y.P., Bransteitter, R., Chelico, L., Sen, U., Stevens, R.C., Goodman, M.F. and Chen, X.S. (2008) Crystal structure of the anti-viral APOBEC3G catalytic domain and functional implications. Nature, 456, 121–124.

34. Chelico, L., Sacho, E.J., Erie, D.A. and Goodman, M.F. (2008) A model for oligomeric regulation of APOBEC3G cytosine deaminase-dependent restriction of HIV. J. Biol. Chem., 283, 13780–13791.

35. Furukawa, A., Nagata, T., Matsugami, A., Habu, Y., Sugiyama, R., Hayashi, F., Kobayashi, N., Yokoyama, S., Takaku, H. and Katahira, M. (2009) Structure, interaction and real-time monitoring of the enzymatic reaction of wild-type APOBEC3G. EMBOJ., 28, 440–451.

36. Chelico, L., Prochnow, C., Erie, D.A., Chen, X.S. and Goodman, M.F. (2010) Structural model for deoxycytidine deamination mechanisms of the HIV-1 inactivation enzyme APOBEC3G. J. Biol. Chem., 285, 16195–16205.

37. Kitamura, S., Ode, H., Nakashima, M., Imahashi, M., Naganawa, Y., Kurosawa, T., Yokomaku, Y., Yamane, T., Watanabe, N., Suzuki, A. et al. (2012) The APOBEC3C crystal structure and the interface for HIV-1 Vif binding. Nat. Struct. Mol. Biol., 19, 1005–1010.

38. Siu, K.K., Sultana, A., Azimi, F.C. and Lee, J.E. (2013) Structural determinants of HIV-1 Vif susceptibility and DNA binding in APOBEC3F. Nat Commun, 4, 2593.

39. Byeon, I.J., Ahn, J., Mitra, M., Byeon, C.H., Hercik, K., Hritz, J., Charlton, L.M., Levin, J.G. and Gronenborn, A.M. (2013) NMR structure of human restriction factor APOBEC3A reveals substrate binding and enzyme specificity. Nat Commun, 4, 1890.

40. Mitra, M., Hercik, K., Byeon, I.J., Ahn, J., Hill, S., Hinchee-Rodriguez, K., Singer, D., Byeon, C.H., Charlton, L.M., Nam, G. et al. (2014) Structural determinants of human APOBEC3A enzymatic and nucleic acid binding properties. Nucleic Acids Res., 42, 1095-1110.

41. Lu, X., Zhang, T., Xu, Z., Liu, S., Zhao, B., Lan, W., Wang, C., Ding, J. and Cao, C. (2015) Crystal structure of DNA cytidine deaminase ABOBEC3G catalytic deamination domain suggests a binding mode of full-length enzyme to single-stranded DNA. J. Biol. Chem., 290, 4010–4021.

42. Shi, K., Carpenter, M.A., Kurahashi, K., Harris, R.S. and Aihara, H. (2015) Crystal Structure of the DNA Deaminase APOBEC3B Catalytic Domain. j. Biol. Chem., 290, 28120–28130.

43. Shaban, N.M., Shi, K., Li, M., Aihara, H. and Harris, R.S. (2016) 1.92 Angstrom Zinc-Free APOBEC3F Catalytic Domain Crystal Structure. j. Mol. Biol., 428, 2307–2316.

44. Byeon, I.J., Byeon, C.H., Wu, T., Mitra, M., Singer, D., Levin, J.G. and Gronenborn, A.M. (2016) Nuclear Magnetic Resonance Structure of the APOBEC3B Catalytic Domain: Structural Basis for Substrate Binding and DNA Deaminase Activity. Biochemistry, 55, 2944–2959.

45. Xiao, X., Li, S.X., Yang, H. and Chen, X.S. (2016) Crystal structures of APOBEC3G N-domain alone and its complex with DNA. Nat Commun, 7, 12193.

46. Kouno, T., Silvas, T.V., Hilbert, B.J., Shandilya, S.M.D., Bohn, M.F., Kelch, B.A., Royer, W.E., Somasundaran, M., Kurt Yilmaz, N., Matsuo, H. et al. (2017) Crystal structure of APOBEC3A bound to single-stranded DNA reveals structural basis for cytidine deamination and specificity. Nat Commun, 8, 15024.

47. Shi, K., Carpenter, M.A., Banerjee, S., Shaban, N.M., Kurahashi, K., Salamango, D.J., McCann, J.L., Starrett, G.J., Duffy, J.V., Demir, O. et al. (2017) Structural basis for targeted DNA cytosine deamination and mutagenesis by APOBEC3A and APOBEC3B. Nat. Struct. Mol. Biol., 24, 131–139.

48. Bowers, K.J., Chow, E., Xu, H., Dror, R.O., Eastwood, M.P., Gregersen, B.A., Klepeis, J.L., Kolossvary, I., Moraes, M.A., Sacerdoti, F.D., Salmon, J.K., Shan, Y. and Shaw, D.E. (2006), Proceedings of the 2006 ACM/IEEE conference on Supercomputing, pp. 84.

49. Harder, E., Damm, W., Maple, J., Wu, C., Reboul, M., Xiang, J.Y., Wang, L., Lupyan, D., Dahlgren, M.K., Knight, J.L. et al. (2016) OPLS3: A Force Field Providing Broad Coverage of Drug-like Small Molecules and Proteins. j. Chem. Theory Comput., 12, 281–296.

50. Leidner, F., Kurt Yilmaz, N., Paulsen, J., Muller, Y.A. and Schiffer, C.A. (2018) Hydration Structure and Dynamics of Inhibitor-Bound HIV-1 Protease. j. Chem. Theory Comput., In press.

51. Flyvbjerg, H. and Petersen, H.G. (1989) Error estimates on averages of correlated data. J. Chem. Phys., 91, 461–466.

52. Shi, K., Demir, O., Carpenter, M.A., Wagner, J., Kurahashi, K., Harris, R.S., Amaro, R.E. and Aihara, H. (2017) Conformational Switch Regulates the DNA Cytosine Deaminase Activity of Human APOBEC3B. Sci. Rep., 7, 17415.

53. Rathore, A., Carpenter, M.A., Demir, O., Ikeda, T., Li, M., Shaban, N.M., Law, E.K., Anokhin, D., Brown, W.L., Amaro, R.E. et al. (2013) The local dinucleotide preference of APOBEC3G can be altered from 5’-CC to 5’-TC by a single amino acid substitution. J. Mol. Biol., 425, 4442–4454.

54. Sharma, S., Patnaik, S.K., Taggart, R.T., Kannisto, E.D., Enriquez, S.M., Gollnick, P. and Baysal, B.E. (2015) APOBEC3A cytidine deaminase induces RNA editing in monocytes and macrophages. Nat Commun, 6, 6881.

